# The potential of ensemble-based automated sleep staging on single-channel EEG signal from a wearable device

**DOI:** 10.1101/2025.07.21.665164

**Authors:** F. Salfi, D. Corigliano, G. Amicucci, S. Mombelli, A. D’Atri, J. Axelsson, M. Ferrara

## Abstract

Machine-learning-based sleep staging models have achieved expert-level performance on standard polysomnographic (PSG) data. However, their application to EEG recorded by wearable devices remains limited by non-conventional referencing montage and the lack of benchmarking against PSG. Here, we tested whether an ensemble of state-of-the-art automatic staging algorithms can reliably classify sleep from a customized configuration of the ZMax headband, adapted to record a single fronto-mastoid EEG channel.

A total of 35 nights of simultaneous ZMax and PSG recordings were acquired in a home setting, amounting to 250.02 hours of analysable data from 10 healthy participants. PSG data were scored according to AASM criteria by two independent experts from different sleep centres, with discrepancies resolved to obtain a consensus hypnogram. ZMax signal was processed using four machine-learning algorithms (YASA, U-Sleep, SleepTransformer, DeepResNet), whose predictions were further combined into a final ensemble scoring through *soft-voting*.

The ensemble scoring achieved almost perfect agreement with human consensus staging (night-level mean ± SD; accuracy = 88.83% ± 2.84%, Cohen’s κ = 84.10% ± 4.52%, and Matthews Correlation Coefficient = 84.54% ± 4.23%). It showed excellent predictive accuracy for REM (F1-score = 93.99%), N3 (89.53%), N2 (87.93%), and wakefulness (86.37%), with lower performance for N1 (53.20%).

These findings support the deployment of an ensemble scoring approach based on state-of-the-art sleep staging algorithms on ultra-minimal, mastoid-referenced EEG setups. This paradigm opens the way to the integration of data from modern wearable technologies into traditional PSG-based sleep research, overcoming longstanding barriers to ecological and large-scale sleep monitoring.

## Introduction

Polysomnography (PSG) remains the accepted *gold standard* for objective evaluation of human sleep. A full-night PSG typically consists of the simultaneous measurement of cortical electroencephalogram (EEG) channels together with electro-oculography (EOG), sub-mental electromyography (EMG), electrocardiography, airflow, respiratory effort, pulse oximetry, and body position. Following acquisition, sleep stages are manually assigned by a trained scorer to consecutive 30-second epochs based on characteristic pamerns observed in EEG, EOG, and EMG channels according to the American Academy of Sleep Medicine (AASM) scoring rules^1^, producing a hypnogram that forms the basis for clinical and research inferences.

Despite its unrivalled information content, routine PSG suffers from important practical constraints. Laboratory studies require expensive equipment and overnight technical supervision, meaning that only a small portion of the population will ever receive an objective assessment of their sleep across the lifespan, typically limited to diagnostic evaluations for suspected sleep disorders. Moreover, the unfamiliar laboratory environment can itself alter sleep architecture and quality, especially during the first night in the novel sleep seming (the well-documented *first-night effect*^*2*^), leading recordings to not fully reflect habitual sleep. Manual sleep staging, while designed to maximise comparability across centres, adds another layer of complexity. An experienced technologist typically spends 1–2 h per record, yet inter-rater agreement across laboratories still hovers in the “moderate-to-substantial” range (Cohen’s κ ≈ 0.60–0.80)^3–6^. This variability reduces statistical power, undermines the generalizability and reproducibility of findings, and complicates multi-site clinical trials.

Technological advances over the past decade offer plausible remedies for both bomlenecks. Lightweight wearable EEG systems (headbands, ear-EEG devices and other form factors) now deliver signal quality approaching that of laboratorial amplifiers while enabling truly ecological, self-applied recordings in the sleeper’s own bedroom^7–9^. Recent feasibility studies demonstrate high user-tolerance and good night-to-night compliance^9^, fuelling rapid market growth and research interest in this sector. In parallel, machine-learning–based sleep scoring algorithms have reached high levels of accuracy, equalling^10^ or even surpassing^11^ the human inter-scorer agreement, opening the way for fully automated, accessible, and reproducible sleep staging. A recent innovation in this context is the ensemble scoring approach, in which multiple automatic sleep staging systems are applied in parallel to the same EEG signal, and their outputs are integrated through a probabilistic combination scheme. A recent benchmarking study^11^ demonstrated that the ensemble-based method outperformed each of their individual component algorithms across various sleep stages and large-scale datasets (e.g., those from the National Sleep Research Resource - NSRR^12^), standing as a potential best-practice solution for automated sleep staging.

Intriguingly, the same investigation^11^ described the release of pre-trained models based on high-performing architectures to support a single-channel input format, thereby promoting their implementation in simplified sleep-monitoring setups. However, the non-conventional referencing used by many wearable EEG devices—often replacing a mastoid reference with closely spaced frontal electrodes to enhance usability—compromises the applicability of state-of-the-art algorithms trained on standard PSG signals. Indeed, many manufacturers and developers of wearable EEG systems have designed their own proprietary staging algorithms, tailored and validated specifically for use with the unique signal characteristics of their device. While device-specific solutions may achieve good performance, they pose significant limitation for scientific generalizability by preventing direct comparability between studies using different hardware, including conventional PSG, and thus contribute to fragmentation in the basic and clinical sleep research field.

In this proof-of-concept study, we tested a pragmatic solution to enable the application of an ensemble scoring approach^11^ based on state-of-the-art sleep staging algorithms^13–16^ to a single-channel EEG signal acquired from a commercially available wearable headband. Specifically, we replaced the default frontal reference of the widely distributed ZMax device (Hypnodyne Corp., Bulgaria) with a contralateral mastoid electrode, resulting in a fronto-mastoid derivation analogous to the standard AF8–M1 channel. We then evaluated the performance of the ensemble-based automated scoring paradigm when applied over 35 nights of ZMax recordings by comparing the resulting hypnograms and macrostructure metrics with those obtained from the consensus of two experienced sleep experts who independently scored co-recorded PSG data.

We hypothesised that restoring a PSG-compatible signal configuration within the ZMax device could enable full exploitation of the potential of existing PSG-trained autoscoring models. In particular, combining an ensemble-based classification strategy with a single-channel, mastoid-referenced EEG setup may offer a scalable and user-friendly solution for reliable home sleep monitoring, while preserving direct comparability with the gold standard of sleep research.

## Methods

### Sleep EEG acquisition

A total of 42 nights of simultaneous home-based sleep EEG co-recordings were collected from 10 participants (mean age ± standard deviation (SD); 26.10 ± 3.78, 6 females) using both a wearable EEG-based system (ZMax headband, Hypnodyne Corp., Bulgaria) and a standard PSG amplifier (Micromed SD 32R, Micromed S.p.A., Italy). Participants were included based on a screening interview if they had no known history of psychiatric, neurological, or sleep disorders and were not undergoing any pharmacological treatment that could interfere with sleep architecture. The study was approved by the Multidisciplinary Internal Review Board of the University of L’Aquila (ID number: 03/2025). Due to technical issues affecting ZMax recordings (3 nights) and PSG recordings (4 nights), the final dataset consisted of 35 analysable nights amounting to 266.97 hours of data. Due to signal quality issues, a total of 16.95 hours across 5 of the retained nights were excluded from the analyses, as those portions of data did not meet the minimum criteria for AASM-compliant scoring due to electrode dropouts. The final analysable dataset thus consisted of 250.02 hours, corresponding to 30,002 30-second epochs.

The ZMax headband is a commercially available device designed to record sleep EEG in home settings. It is easy to apply and comfortable to wear, consisting of a small frontal unit connected via four snap connectors to an Ag/AgCl-based biocompatible hydrogel patch. This configuration allows for the acquisition of two fronto-frontal channels (F7–Fpz and F8–Fpz), using a frontal ground. Data are acquired with a 0.1-128 Hz bandwidth, sampled at 256 Hz and stored locally on an internal microSD card.

To meet the aims of the present study, we adapted standard laboratory electrode cables to make them compatible with the ZMax connectors, allowing full control over electrode placement. This modification enabled us to shift the reference from a frontal to a mastoid site, allowing frontal electrodes (AF7 and AF8) to be referenced to M1. Electrodes were disposable, self-adhesive Ag/AgCl snap-type electrodes (Kendall ARBO, Ø24 mm, solid hydrogel), connected using snap-lead wires (MedCat, Netherlands), ensuring fast application, stable contact throughout the night, and high user comfort.

For the PSG recordings, we used a minimal AASM-compliant montage consisting of three cortical EEG channels (F4–M1, C4–M1, O2–M1), bilateral EOG, and sub-mental EMG. EEG signals were recorded using Ag/AgCl cup electrodes applied with EC2+ Grass Electrode Cream (Natus Neurology, United States). PSG data were sampled at 256 Hz and stored applying the filtering AASM recommendations (0.3–35 Hz band-pass filtering for EEG and EOG; 10–100 Hz for EMG)^1^.

All electrodes were applied, and both recording systems were fully set up, by trained researchers (F.S, D.C., and G.A.) during evening home visits. Participants were instructed on how to start and end the recording procedures. At “lights-off” time participants were asked to perform five consecutive horizontal eye movements and five brief teeth clenches, which were subsequently used for synchronization of the ZMax and PSG traces.

### Manual sleep scoring

ZMax and PSG files were synchronised visually using the predefined eye movements performed by participants at the “lights-off’’ time. Two experienced sleep technologists from independent sleep centres (D.C. and S.M.) then performed manual sleep staging independently on PSG data following the AASM scoring rules^1^, each blind to the other’s scoring and to the wearable data. Disagreements were resolved through joint review of each mismatched epoch to obtain a consensus hypnogram, which served as the ground-truth reference.

### Automatic sleep scoring

The raw AF8-M1 ZMax signal was analysed using SLEEPYLAND^11^, an open-source web-based platform for automated sleep staging that integrates multiple state-of-the-art deep learning and feature-based algorithms. Specifically, ZMax data were submitted to four distinct classifiers: Yet another Spindle Algorithm (YASA)^16^, U-Sleep^15^, SleepTransformer^13^, and DeepResNet^14^. Each of these algorithms was developed adopting complementary modeling strategies, spanning handcrafted features, convolutional and recurrent networks, and transformer-based sequence encoders. All models demonstrated high performance on large, publicly available PSG datasets^13–16^.

YASA is a feature-based algorithm employing gradient-boosted decision trees trained on 3,163 full-night PSG recordings (over 30,000 hours) across heterogeneous populations. Validation study^16^ demonstrated a median night-level accuracy of 87.56% and a Cohen’s κ of 81.88% across diverse test sets. U-Sleep is a convolutional neural network inspired by the U-Net architecture. It was trained and evaluated on 19,924 PSG recordings from 15,660 subjects from 16 clinical studies and has been shown to match the performance the of best human expert reaching a mean F1 score of 79%^15^. SleepTransformer is a transformer-based sequence-to-sequence model. Validation study on PSGs from the Sleep Heart Health Study (n = 5,791) and the 2018 version of the Sleep-EDF Expanded dataset (n = 79) reported performance comparable to other state-of-the-art systems (accuracy up to 87.7%, Cohen’s κ of 82.8%)^13^. Finally, DeepResNet is a residual convolutional neural network conceived as an adaptation of a ResNet architecture. The model was developed on 15,684 PSGs, achieving an accuracy of 86.90% and a Cohen’s κ of 79.90%^14^ in a training configuration including all the available data sources^14^.

While the original implementations of these models were often constrained to specific signal configurations, SLEEPYLAND allows inference on any derivation and includes retrained versions of U-Sleep, SleepTransformer, and DeepResNet to support flexible input formats, including the single-channel EEG configuration used in this study. Specifically, sleep staging algorithms were re-trained on all available NSRR datasets, amounting to ∼220,000 hours of PSG data from 24 clinical cohorts. Technical details on model training and evaluation procedures can be found in the original SLEEPYLAND documentation^11^.

Each classifier produced a 30-second resolution hypnogram, along with a probability distribution indicating the likelihood assigned to each individual sleep stage. These probability scores formed the basis for the final stage classification returned by the algorithm. Intriguingly, the SLEEPYLAND platform also enables ensemble scoring by combining the outputs of multiple models based on *soft voting* approach. Specifically, it computes the average probability for each sleep stage across all algorithms and assigns to each epoch the stage with the highest mean probability. This ensemble approach leverages the complementary strengths of different models while mitigating individual biases and limitations, ultimately generating a consensus hypnogram that is more robust and reliable than the output of any single model alone^11^.

### Evaluation metrics and analyses

The agreement between manual sleep staging (i.e., the ground-truth) and the automatic scoring of four algorithms (YASA^16^, U-Sleep^15^, SleepTransformer^13^, DeepResNet^14^) and a final ensemble model was assessed computing four complementary metrics. *Accuracy* reflects the overall proportion of correctly classified epochs, regardless of sleep stage. *Cohen’s κ coefficient*^17^ adjusts for chance agreement and provides a robust estimate of inter-rater reliability, particularly in cases of class imbalance. *Matthews correlation coefficient*^18^ (MCC) is a balanced measure that takes into account true and false positives and negatives across all classes, generally regarded as one of the most informative metrics for evaluating classification performance, especially in the presence of class imbalance. Moreover, we computed *F1 scores* for each sleep stage as the harmonic mean of precision and recall. This metric is particularly informative in multi-class classification contexts, as it balances false positives and false negatives, and offers a comprehensive view of per-stage classification performance. We also generated confusion matrices for each method to provide a detailed overview of classification behaviour. Matrices reports, for each true sleep stage, the percentage of epochs predicted in each category, thereby highlighting stage-specific classification performance. The diagonal elements represent recall (also referred to as sensitivity), that is, the proportion of epochs of a given stage correctly identified by the model. All metrics were estimated at single-night level and averaged across nights, ensuring that each sleep recording contributed equally to the final group-level performance estimates and allowing for the assessment of inter-night variability across models. Pooled epoch-level metrics are reported in the Supplementary Material.

Furthermore, we evaluated how each automatic method replicated sleep macrostructure as estimated by manual scoring. These analyses were conducted after excluding the 5 nights that contained missing scoring epochs due to signal quality issues. For each night and scoring system, we computed classical sleep parameters including total sleep time (TST), sleep efficiency (SE%), wake after sleep onset (WASO), and sleep onset latency (SOL). We also calculated the relative (%) and absolute (min) time spent in each sleep stage (N1, N2, N3, REM).

To statistically assess the differences between each automatic method (including the ensemble model) and the manual consensus, we conducted two-sided paired t-tests on each sleep parameter. The resulting p-values were adjusted for multiple comparisons using the Holm method. In addition, Bland–Altman plots comparing the ensemble model to the human consensus were generated to visually assess agreement, identify potential systematic biases, and evaluate the dispersion of differences across the range of values for each macrostructural parameter.

Finally, to accommodate scenarios in which distinguishing N1 from N2 may be less critical, we also repeated all the above-described analyses using a simplified four-class scheme in which N1 and N2 were combined into a single “Light sleep” stage.

All parameters and metrics were computed using custom Python scripts based on functions from *scikit-learn*, and visualizations were created with *seaborn* and *matplotlib*. Statistical analyses were performed using *R software* (version 4.1.2; R Foundation for Statistical Computing, Austria).

## Results

Table 1 reports the performance metrics averaged across nights for each automatic scoring model. The ensemble approach achieved *almost perfect* agreement with human consensus staging, as classified according with the original interpretation guidelines for Cohen’s κ^19^. Among the individual algorithms, DeepResNet showed the best agreement, while all deep learning-based models outperformed the feature-based YASA algorithm. Stage N1 was consistently the most challenging to classify across models.

**Table 1.** Performance metrics (inter-night mean percentage ± standard deviation) for each automatic sleep staging algorithm and the ensembled model across all nights, computed with respect to the consensus of two human scorers.

| Metric | YASA | U-Sleep | SleepTransformer | DeepResNet | Ensemble |
| --- | --- | --- | --- | --- | --- |
| Accuracy (%) | 84.26 ± 3.16 | 87.03 ± 3.27 | 87.61 ± 2.54 | 88.30 ± 2.59 | 88.83 ± 2.84 |
| Cohen's $\kappa$ (%) | 77.86 ± 4.92 | 81.51 ± 5.09 | 82.47 ± 3.85 | 83.39 ± 3.97 | 84.10 ± 4.52 |
| MCC (%) | 78.30 ± 4.59 | 82.22 ± 4.56 | 82.82 ± 3.65 | 83.77 ± 3.77 | 84.54 ± 4.23 |
| F1 N1 (%) | 34.15 ± 11.50 | 52.33 ± 13.01 | 49.00 ± 12.13 | 54.87 ± 11.98 | 53.20 ± 13.56 |
| F1 N2 (%) | 84.14 ± 4.98 | 86.12 ± 4.80 | 86.51 ± 3.53 | 87.44 ± 3.48 | 87.93 ± 4.13 |
| F1 N3 (%) | 89.35 ± 4.12 | 85.74 ± 6.94 | 88.81 ± 4.89 | 88.79 ± 5.34 | 89.53 ± 5.06 |
| F1 REM (%) | 87.60 ± 7.61 | 93.63 ± 5.05 | 93.22 ± 4.82 | 93.84 ± 3.96 | 93.99 ± 5.16 |
| F1 Wake (%) | 75.54 ± 11.81 | 85.25 ± 8.55 | 85.32 ± 7.75 | 85.20 ± 8.94 | 86.37 ± 8.62 |

Abbreviations: MCC, Matthews correlation coefficient.

The ensemble model performed even better when applied to the four-stage classification scheme, achieving an accuracy of 91.01 ± 2.89%, a Cohen’s κ of 86.63 ± 4.60%, and an MCC of 87.00 ± 4.28%. The F1-score for the Light sleep category reached 89.92 ± 3.66%.

Further examination of the confusion matrix (Figure 1) showed high sensitivity of all models for each sleep stage, except for N1 sleep. The two most common inaccuracies of both the algorithms and the ensemble model were mislabeling N1 stage as N2 or Wake, and mislabeling N3 as N2 sleep.

**Figure 1.**
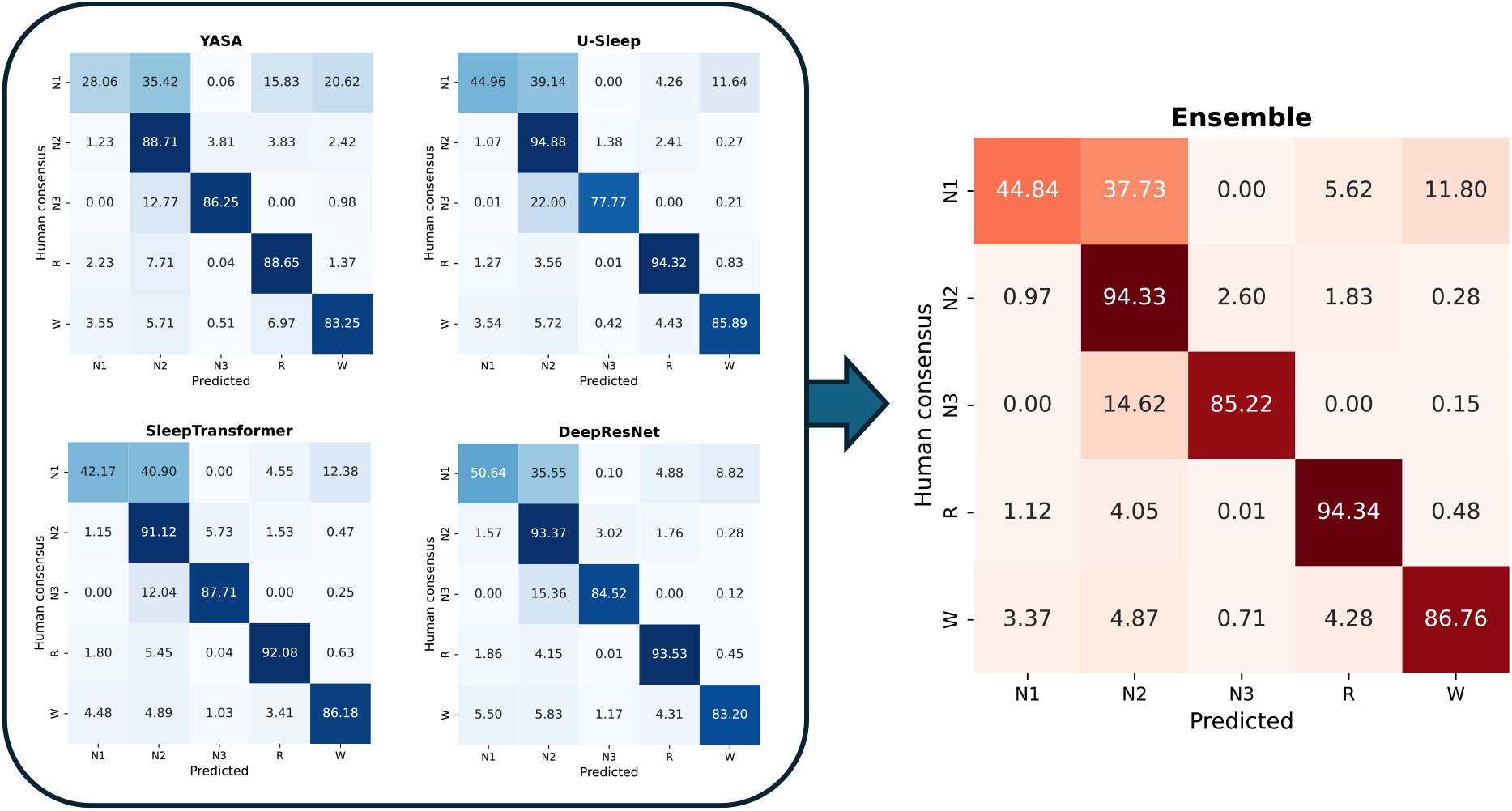
Confusion matrices for each automatic sleep staging algorithm (blue colormaps) and the ensemble model (red colormap), averaged across nights. Diagonal elements represent the percentage of epochs correctly classified for each sleep stage (i.e., sensitivity or recall) with respect to the consensus of two human scorers, while off-diagonal elements indicate misclassifications.

Pooled epoch-level analyses returned almost identical metrics and results (see Table S1 and Figure S1).

Table 2 summarizes the key sleep metrics derived from each automatic scoring algorithm and the human consensus. Sleep macrostructure estimates obtained from the ensemble model showed high concordance with those derived from manual scoring. Core metrics such as TST, SE%, WASO, and SOL were closely matched between the two scoring systems, with no significant deviations observed. Stage-level comparisons revealed the most pronounced differences in the classification of N1 and N3 sleep, where the ensemble model slightly underestimated both stages. These discrepancies were counterbalanced by a tendency to overestimate N2 sleep. Finally, REM sleep estimates remained consistent with the ground truth. Bland–Altman plots (Figure 2) confirmed optimal concordance across most sleep parameters, with the vast majority of data points falling within the 95% limits of agreement and no evident proportional bias.

**Table 2.** Sleep macrostructure metrics estimated by automatic sleep staging models and human consensus.

| Variable | YASA | U-Sleep | SleepTransformer | DeepResNet | Ensemble | Human Consensus |
| --- | --- | --- | --- | --- | --- | --- |
| TST (min) | 418.13 ± 57.69 | 424.32 ± 59.51 | 423.93 ± 59.95 | 426.25 ± 59.71 | 424.33 ± 59.46 | 424.30 ± 58.28 |
| SE (%) | 90.60 ± 4.09* | 91.92 ± 4.05 | 91.83 ± 4.23 | 92.35 ± 4.10 | 91.92 ± 4.04 | 91.98 ± 4.12 |
| WASO (min) | 32.18 ± 18.22 | 28.10 ± 17.66 | 28.22 ± 17.54 | 26.42 ± 17.68 | 27.33 ± 17.48 | 28.65 ± 16.14 |
| SOL (min) | 11.27 ± 6.51* | 9.17 ± 7.19 | 9.43 ± 7.54 | 8.92 ± 7.11 | 9.92 ± 7.24 | 8.63 ± 8.11 |
| N1 (%) | 2.78 ± 1.86*** | 3.42 ± 1.40*** | 3.55 ± 1.03*** | 4.24 ± 1.02*** | 3.28 ± 1.07*** | 5.23 ± 1.06 |
| N2 (%) | 44.44 ± 5.99*** | 48.18 ± 6.13*** | 44.73 ± 5.49*** | 45.74 ± 5.88*** | 45.98 ± 5.61*** | 40.82 ± 4.87 |
| N3 (%) | 24.79 ± 5.20* | 20.81 ± 5.09*** | 25.30 ± 4.69 | 23.10 ± 5.04** | 23.37 ± 4.83** | 26.42 ± 3.91 |
| REM (%) | 27.98 ± 3.51 | 27.58 ± 3.15 | 26.42 ± 3.22 | 26.92 ± 3.43 | 27.37 ± 3.10 | 27.54 ± 3.69 |
| N1 (min) | 11.60 ± 7.24*** | 14.32 ± 5.38*** | 14.77 ± 4.21*** | 17.75 ± 3.78*** | 13.72 ± 4.49*** | 21.80 ± 4.84 |
| N2 (min) | 186.28 ± 37.42** | 205.08 ± 40.11*** | 189.75 ± 37.89*** | 195.47 ± 37.13*** | 195.45 ± 40.52*** | 173.53 ± 35.77 |
| N3 (min) | 103.40 ± 26.91** | 87.58 ± 22.99*** | 107.10 ± 21.69 | 98.00 ± 25.47** | 98.70 ± 20.51** | 112.15 ± 20.08 |
| REM (min) | 116.85 ± 21.51 | 117.33 ± 22.35 | 112.32 ± 22.90 | 115.03 ± 22.86 | 116.47 ± 22.50 | 116.82 ± 21.69 |

*Abbreviations:* TST, total sleep time; SE, sleep efficiency; WASO, wake after sleep onset; SOL, sleep onset latency.
*Notes:* Asterisks indicate statistically significant differences (\* $p < 0.05$ , \*\* $p < 0.01$ , \*\*\* $p < 0.001$ ; row-wise adjusted for multiple comparisons using the Holm method) between each autoscoring system and the human consensus.

**Figure 2.**
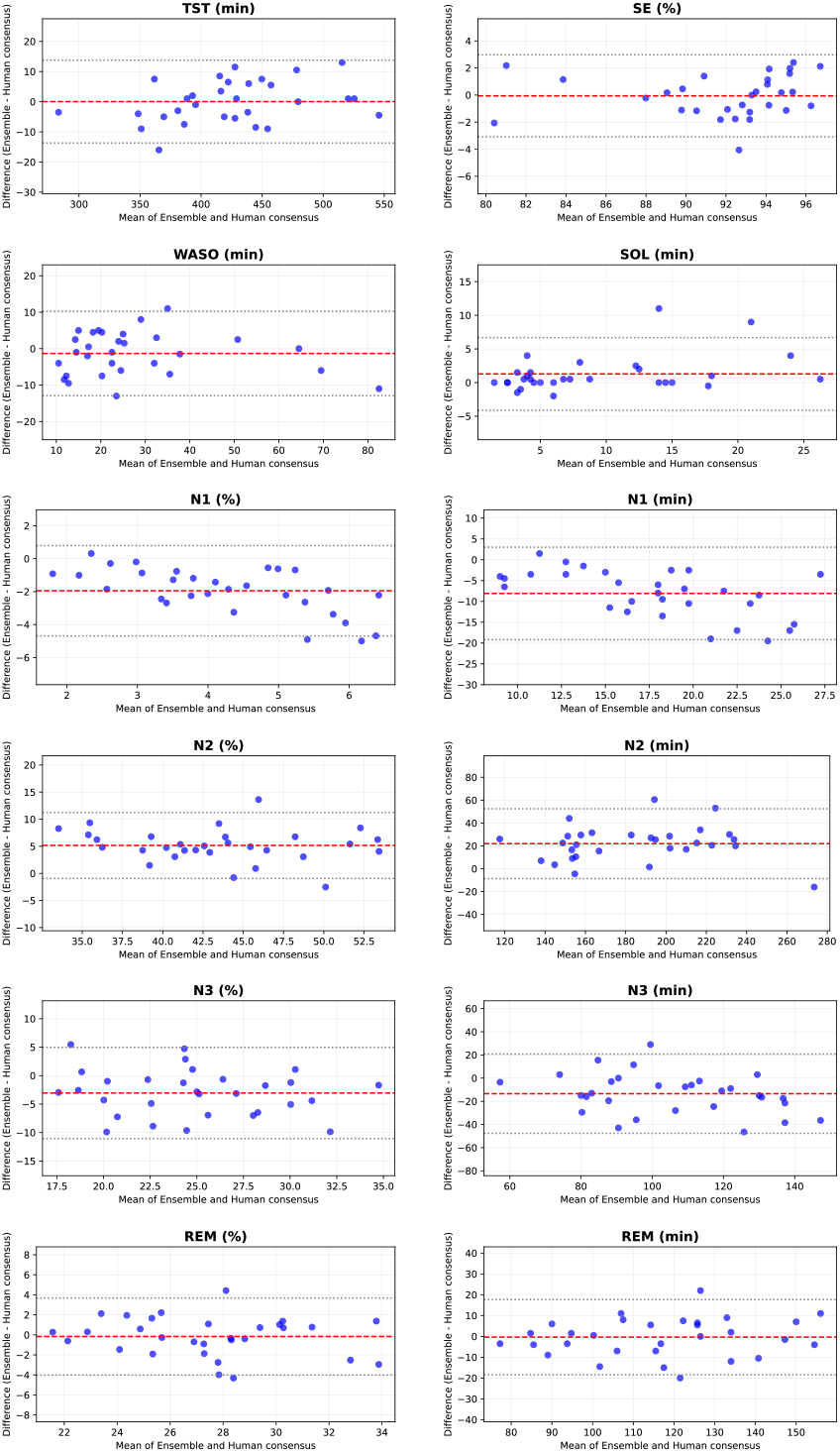
Bland–Altman plots comparing macrostructural sleep parameters derived from the ensemble model and the human consensus staging. The red dashed line indicates the average difference, while the dotted grey lines mark the 95% limits of agreement (±1.96 SD).

## Discussion

This study evaluated the feasibility of applying an ensemble of state-of-the-art automatic sleep staging algorithms to a single-channel, mastoid-referenced EEG signal obtained using a customised configuration of the ZMax wearable device. Our findings demonstrate that this minimal EEG setup can match classification performance of trained human scorers, particularly when an ensemble method is used.

Across all metrics, the ensemble model outperformed its individual components, achieving a mean accuracy of 88.83% and a Cohen’s κ of 84.10%, falling within or even surpassing the range of inter-rater agreement typically reported in manual PSG staging^3,5^. These findings align with previous work introducing SLEEPYLAND^11^ that demonstrated how an ensemble approach improved sleep staging accuracy compared to individual scoring algorithms in 94.9% of cases. Therefore, our study supported that combining predictions from multiple models enhances classification performance by integrating complementary strengths and mitigating individual model biases. Class-wise F1 scores further confirmed the ensemble’s robustness, with optimal performance across all sleep stages (range: 86.37%–93.99%), except for N1 sleep (53.20%), which remains a known source of ambiguity even among trained scorers^5^.

Regarding sleep macrostructure, the ensemble model accurately replicated key parameters commonly derived from hypnograms, including TST, SE%, WASO, and SOL. However, while estimates for REM sleep were closely aligned with the human consensus (−0.17%, −0.35 min), stage-specific biases emerged for other sleep stages. In particular, the model tended to underestimate N1 (−1.95%, −8.08 min) and N3 (−3.05%, −13.45 min), while overestimating N2 (+5.16%, +21.92 min). This pattern mirrors the most common misclassification trends observed, with both individual models and the ensemble occasionally mislabelling N1 and N3 as N2.

When examining the performance of individual classifiers, it is evident that all deep learning models outperformed the feature-based YASA algorithm. However, it is important to note that among the four algorithms integrated in the SLEEPYLAND platform, YASA is the only one that was not retrained on a single-channel configuration using the full NSRR dataset. Thus, differences in performance should be considered in the context of model retraining procedures and input configurations, which were not harmonized across algorithms. Despite this limitation, our results closely mirror those reported in the original validation study^16^, where YASA achieved a median accuracy of 85.12% and a median Cohen’s κ of 0.78 when evaluated on single-channel EEG input.

A recent validation study^20^ analysed 135 nights of simultaneous recording between PSG and ZMax, aiming to evaluate the performance of ZLab (the proprietary autoscoring system) alongside a novel algorithm (DreamentoScorer) specifically developed to classify sleep stages from the traditional fronto-frontal ZMax signal. The authors demonstrated that DreamentoScorer outperformed ZLab, achieving an overall accuracy of 79.67% and a Cohen’s κ of 72.18% when benchmarked against human scoring. Compared to these results, the approach proposed in our investigation yielded a consistent improvement, both in terms of overall agreement (accuracy = +9.16%, Cohen’s κ = +11.92%) and stage-wise classification performance (F1-score N1 = +13.47%, N2 = +6.70%, N3 = +7.25%, REM = +10.98%, Wake = +9.01%). Overall, these findings suggest that a model-agnostic framework can achieve higher classification performance than device-specific solutions, with the added benefit of interoperability across different systems and research settings. However, it should be noted that the comparison with DreamentoScorer remains indirect, as the present study did not include a head-to-head evaluation using the same dataset or experimental conditions.

The proposed approach opens new avenues for the use of ZMax in advanced applications targeting microstructural EEG features. Visual inspection of the mastoid-referenced signal revealed a striking overlap with the corresponding PSG derivation (F4–M1), suggesting that the signal quality can support the reliable detection of discrete graphoelements such as, for example, sleep spindles and slow oscillations. An additional implication of the customized referencing scheme relates to the deployment of minimal, single-channel EEG systems in non-laboratory environments for advanced neurotechnological applications such as phase-targeted neuromodulation. When adapted with a physiologically neutral reference, systems like ZMax overcome a fundamental limitation of many consumer-grade EEG devices that rely on fronto-frontal montages. These unconventional configurations preclude meaningful polarity interpretation, a prerequisite for implementing phase-targeted auditory stimulation (PTAS)—also known as closed-loop auditory stimulation^21^—and other phase-dependent protocols requiring precise identification of the slow oscillation phase like closed-loop targeted memory reactivation^22,23^. This rationale has directly motivated our effort to customize the ZMax referencing scheme, resulting in promising preliminary outcomes for the application of PTAS in detecting and modulating slow oscillation dynamics in a home setting, with minimal intrusiveness and high ease of use for participants (Salfi et al., in preparation).

It is worth noting that, although in the present study electrode placement for ZMax was performed by trained researchers to ensure maximal comparability with PSG recordings, the proposed novel configuration (based on disposable, self-adhesive snap-type electrodes) only minimally increases setup complexity compared to standard use and does not compromise the device suitability for self-administration by end users, as confirmed by parallel ongoing investigations.

Despite the promising results, some limitations must be acknowledged. This study involved a limited number of participants, which constrains the generalizability of the findings. Moreover, the sample was restricted to healthy young adults without diagnosed sleep disorders. Future research should aim to replicate and extend these findings in clinical populations, as well as evaluate the robustness of the proposed approach across a broader range of demographic and sleep-related characteristics.

In our study, no nights were interrupted due to participant drop-out, and no substantial complaints due to the recording setting were reported. However, we acknowledge that collecting structured data on user experience and technological acceptance would have strengthened our evaluation. Notably, a previous investigation^20^ has already addressed this aspect, demonstrating that the native ZMax configuration is well-tolerated, with no reported impact on sleep quality or cumulative negative effects over consecutive nights. We may hypothesise that similar outcomes would apply in our setting as well.

Finally, the proposed montage required technical adaptation not yet supported in the native ZMax platform. Broader adoption of this strategy will depend on manufacturer integration or standardized adaptors for field use.

## Conclusion

Our findings demonstrate that an ultra-minimal wearable EEG device, when properly adapted with a physiological reference, can support accurate, automated sleep staging using existing PSG-trained algorithms. This approach eliminates the need for model retraining, effectively addressing the comparability issues that have long hindered the integration of data from modern wearable technologies into traditional PSG-based sleep research. Moreover, it highlights the value of ensemble methods in enhancing performance, particularly in ecological settings where data quality may vary. This paradigm may pave the way for large-scale sleep research conducted entirely in naturalistic home environments, unlocking new opportunities for longitudinal and population-level studies.

## Author contributions

**Federico Salfi:** conceptualization, software, data curation, visualization, methodology, investigation, formal analysis, writing – original draft, writing – review and editing. **Domenico Corigliano:** conceptualization, software, data curation, methodology, investigation, formal analysis, writing – review and editing. **Giulia Amicucci:** software, data curation, investigation, formal analysis, writing – review and editing. **Samantha Mombelli:** formal analysis, writing – review and editing. **Aurora D’Atri:** writing – review and editing. **John Axelsson:** methodology, writing – review and editing. **Michele Ferrara:** supervision, writing – review and editing.

## Data availability statement

The data supporting the findings of this study are available from the corresponding author upon reasonable request.

