## Supplementary material for "The potential of ensemble-based automated sleep staging on single-channel EEG signal from a wearable device"

### *Running title:* Automated sleep staging and wearable EEG

Salfi F.[1*†], Corigliano D.[1,2†], Amicucci G.[1], Mombelli S.[3], D'Atri A.[1], Axelsson J.[4,5], and Ferrara M.[1*]

[1]*Department of Biotechnological and Applied Clinical Sciences, University of L'Aquila, L'Aquila, Italy*
[2]*Department of Psychology, Sapienza University of Rome, Rome, Italy*
[3]*Center for Advanced Research in Sleep Medicine, Research, center of the Centre intégré universitaire de santé et de services sociaux du Nord de l'Île-de-Montréal (Hôpital du Sacré-Coeur de Montréal), Montreal, Canada*
[4]*Department of Clinical Neuroscience, Karolinska Institutet, Stockholm, Sweden*
[5]*Department of Psychology, Stockholm University, Stockholm, Sweden*

†Share the first authorship
*Corresponding authors

Dr. Federico Salfi, *Ph.D.*
Department of Biotechnological and Applied Clinical Sciences
University of L'Aquila
Via Vetoio
67100 L'Aquila (AQ)
Italy


Prof. Michele Ferrara, *Ph.D.*
Department of Biotechnological and Applied Clinical Sciences
University of L'Aquila
Via Vetoio
67100 L'Aquila (AQ)
Italy


### Supplementary material

**Table S1.** Performance metrics (percentage) for each automatic sleep staging algorithm and the ensembled model across all 30,002 pooled epochs, computed with respect to the consensus of two human scorers.

| Metric | YASA | U-Sleep | SleepTransformer | DeepResNet | Ensemble |
|---|---|---|---|---|---|
| Accuracy (%) | 84.36 | 86.98 | 87.6 | 88.16 | 88.82 |
| Cohen's κ (%) | 78.36 | 81.84 | 82.81 | 83.56 | 84.45 |
| MCC (%) | 78.48 | 82.32 | 82.91 | 83.75 | 84.67 |
| F1 N1 (%) | 34.72 | 53.47 | 49.42 | 55.71 | 53.94 |
| F1 N2 (%) | 84.7 | 86.56 | 86.99 | 87.7 | 88.37 |
| F1 N3 (%) | 89.23 | 85.59 | 88.91 | 88.75 | 89.46 |
| F1 REM (%) | 88.83 | 94.17 | 93.64 | 94.15 | 94.56 |
| F1 Wake (%) | 77.78 | 87.32 | 87.23 | 87.76 | 88.34 |

*Abbreviations: MCC, Matthews correlation coefficient.*

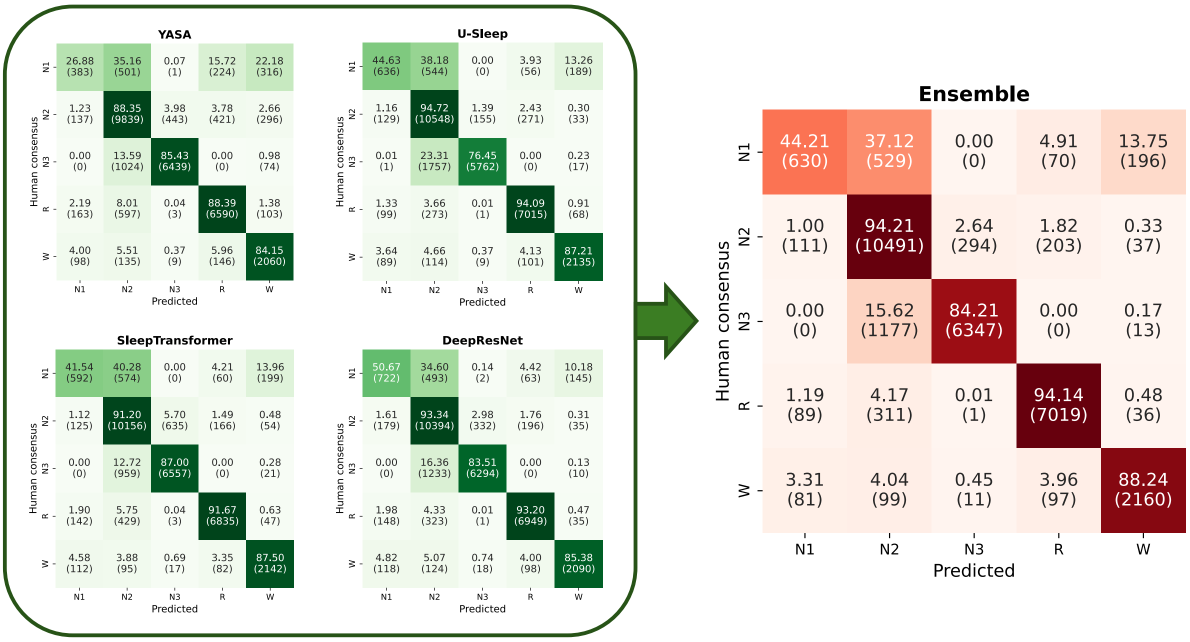


**Figure S1.** Confusion matrices for each automatic sleep staging algorithm (green colormaps) and the ensemble model (red colormap) at the pooled-epoch level. Diagonal elements indicate the percentage of correctly classified epochs for each sleep stage (i.e., sensitivity or recall) compared to the consensus of two expert human scorers, while off-diagonal elements represent misclassifications. The raw number of epochs per cell is reported in parentheses.
